# Explaining temporally clustered errors with an autocorrelated Drift Diffusion Model

**DOI:** 10.64898/2026.03.20.713186

**Authors:** Robin Vloeberghs, Francis Tuerlinckx, Anne E. Urai, Kobe Desender

**Author notes:** **Corresponding author:** Robin Vloeberghs, Brain and Cognition, KU Leuven, Tiensestraat 102, 3000 Leuven, Belgium.

## Abstract

A widely used framework for studying the computational mechanisms of decision-making is the Drift Diffusion Model (DDM). To account for fast and slow errors the DDM incorporates across-trial variability in parameters such as the drift rate and the starting point. Although these variability parameters enable the model to reproduce both fast and slow errors, they rely on the assumption that parameters are independently sampled over trials. As a result, the DDM effectively predicts that errors—whether fast or slow—occur randomly in time. Here, we show that in empirical data this assumption is violated, as error responses are temporally clustered. To address this limitation in the model, we introduce the autocorrelated DDM, in which drift rate, starting point, and boundary evolve from trial-to-trial according to first-order autoregressive (AR1) processes. Using simulations, we demonstrate that, unlike the across-trial variability DDM, the autocorrelated DDM naturally accounts for temporal clustering of errors. We further show that model parameters can be reliably recovered using Amortized Bayesian Inference, even with as few as 500 trials. Finally, in a pre-registered analysis we find that the autocorrelated DDM provides the best account of error clustering across 8 separate datasets, including both human and mouse data. These results establish that accounting for temporal fluctuations in computational parameters is essential for accurately characterizing decision-making behavior.

## Introduction

Every day we make numerous decisions: whether it is safe to cross the street, which t-shirt to wear, or whether to have coffee or tea. To understand the computational mechanisms underlying such decisions, researchers often rely on computational models. Within the context of two-alternative forced choice tasks (2AFC), a well-established and widely used computational model is the Drift Diffusion Model (DDM) (Ratcliff, 1978; Ratcliff & McKoon, 2008; Ratcliff & Tuerlinckx, 2002). Its key tenet is that upon stimulus presentation, the observer accumulates noisy evidence until one of two opposing boundaries is reached (Figure 1A). When a boundary is reached, the response associated with that boundary is selected for execution (e.g., respond ‘left’ or ‘right’). How fast this evidence accumulation reaches the boundary is a function of its slope, quantified by drift rate parameter *v*. With higher values for *v*, the boundary will be reached faster, resulting in shorter and more accurate responses. The amount of evidence that needs to be accumulated is set by the boundary separation parameter *a*. The larger this boundary separation, the smaller the chance of an incorrect response, but at the cost of a longer reaction time (i.e., the speed-accuracy trade-off). The starting point parameter *z* determines where the evidence accumulation process begins and allows to capture *a priori* biases in the decision process, by shifting the starting point closer to one of the two boundaries. Non-decisional processes, such as stimulus encoding and motor execution, are captured by the non-decision time parameter *T*_*er*_.

**Figure 1:**
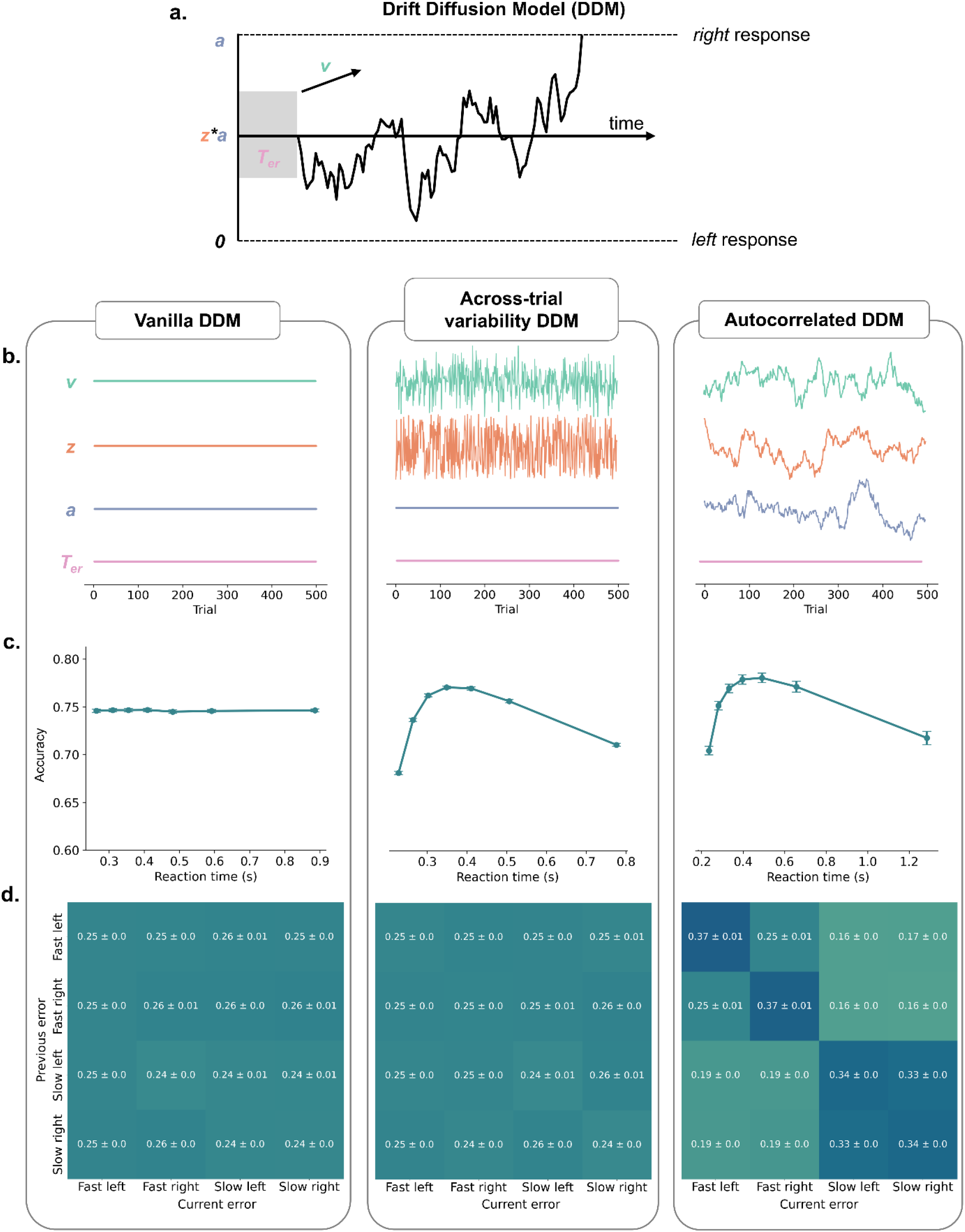
**a**. The drift diffusion model assumes that decisions are made by accumulating noisy evidence over time (governed by drift rate *v*) until one of two opposing boundaries, *a*, are reached. The starting point of the accumulation process is controlled by *z*, and non-decision related aspects are captured by non-decision time, *T*_*er*_. **b**. In the vanilla (four-parameter) version of the model, all parameters are assumed to be constant for all trials (left). In the across-trial variability (six-parameter) version of the model, the drift rate and starting point are drawn on each trial from a noisy IID distribution (middle). In the autocorrelated DDM, the drift rate, starting point, and boundary are assumed to be governed by a first-order autoregressive process (AR1), thus fluctuating from trial-to-trial in an autocorrelated fashion. **c**. A well-known limitation of the vanilla DDM is that it cannot account for the presence of fast and slow errors. This can be appreciated by the fact that it predicts a flat conditional accuracy function (left), whereas adding random noise to the drift rate and starting point creates the inverted-U shaped conditional accuracy pattern typically observed in empirical data (middle). Similarly, this signature also emerges when drift rate, starting point, and boundary fluctuate over trials in an autocorrelated fashion (right). **d**. Going beyond the mere presence of fast and slow errors, the current work proposes a novel key empirical feature: fast and slow errors do not occur randomly but instead are temporally clustered in time. Whereas the vanilla and across-trial variability DDM predict that the presence of fast and slow errors does not depend on whether such an error was committed on the previous trial (left, middle), the autocorrelated DDM does predict such sequential dependencies between error types. For instance, a *fast left* error is more likely to be followed by another *fast left* error (.37) compared to a *slow left* error (.16). A *fast left* error refers to a trial on which the correct response was *right* but and incorrect *left* response was given, and the reaction time was faster than the median reaction time. The repetition matrices are calculated using a subset of trials in which two consecutive responses were errors and show the average conditional probabilities over simulated subjects (± one standard error) for the current error, with .25 representing chance level and the columns summing to 1.

A DDM with these four parameters (*v, a, z, T*_*er*_), hereafter referred to as the *vanilla DDM*, captures several key characteristics of decision-making (e.g., speed-accuracy trade-off, difficulty effects). However, a well-known shortcoming of this four parameter version of the model is that it predicts equal reaction times for correct and error responses. In empirical data, however, this is typically not the case: for some tasks it is well described that error RTs are shorter compared to correct RTs, whereas for other tasks the opposite pattern is observed. To appreciate the presence of fast and slow errors, one can inspect the conditional accuracy function, which visualizes average accuracy as a function of reaction times. A typical dataset from a speeded perceptual decision-making task shows an inverted-U pattern here, with lower accuracy for the fastest RTs (i.e., fast errors) and lower accuracy for the slowest RTs (i.e., slow errors). Contrary to this pattern, the vanilla DDM shows a flat conditional accuracy function (Figure 1B). Historically, this shortcoming in the vanilla DDM is resolved by adding across-trial variability to the drift rate and the starting point (Ratcliff & McKoon, 1999). Specifically, slow errors can be captured within the DDM when the drift rate is not a constant but instead on each trial a drift rate *v*_*t*_ is sampled from a normal distribution around the drift rate *v*_*t*_ ~ *N*(*v, s*_*v*_) (Ratcliff, 1978). Intuitively, on some trials the single-trial drift rate will be very low leading to slow and error-prone responses. Similarly, fast errors can be captured within the DDM when the starting point is not fixed but instead on each trial is sampled from a uniform distribution centered around the starting point *z*_*t*_ ~ *U*(*z* − *s*_*z*_/2, *z* + *s*_*z*_/2) (Laming, 1968). Intuitively, on some trials the starting point will be very close to the incorrect boundary, leading to fast and error-prone responses. When simulating data from the across-trial variability DDM (i.e. with six parameters; *v, a, z, T*_*er*_, *s*_*z*_, *s*_*v*_) the conditional accuracy function indeed shows the inverted-U pattern typically observed in empirical data (Figure 1C). Note, that in this model the single-trial parameters, *v*_1:*T*_ and *z*_1:*T*_, are not explicitly estimated.

In the current work, we investigate the computational underpinnings of a behavioral signature central to errors in human decision-making: the temporal clustering of error responses (Dutilh, van Ravenzwaaij, et al., 2012; Gilden & Wilson, 1995). Whereas the across-trial variability parameters allow the DDM to explain both fast and slow errors, given that single-trial drift rate and starting point are on each trial randomly sampled from a distribution, this version of the model effectively predicts that fast and slow error occur randomly in time. Contrary to this prediction, in the current work we investigate in empirical data and models whether fast and slow error responses follow each other more often than would be expected purely by chance. When simulating data from the across-trial variability parameter DDM, indeed fast and slow errors do not cluster in time but instead randomly occur throughout the data (Figure 1D). This is not surprising, across-trial variability parameters do not predict temporal clustering because they assume *independent and identically distributed* (IID) noise distributions. One way to capture temporally clustered errors within a DDM is to introduce autocorrelated fluctuations in parameters, such as the drift rate or starting point. Autocorrelated fluctuations in the decision boundary may likewise give rise to temporal clustering of errors, as periods of reduced response caution (i.e., a lower decision boundary) increase the likelihood of errors occurring on successive trials (Dutilh, van Ravenzwaaij, et al., 2012). Thus, in the current work, we developed an autocorrelated DDM in which the drift rate, starting point, and boundary evolve over trials according to an AR(1) model. The main feature of an autoregressive model is that it produces autocorrelation, in the current application to the drift rate, starting point, and boundary. As a consequence, trials characterized by high (vs. low) drift rates, biased (vs. unbiased) starting points, and conservative (vs. liberal) boundaries tend to be followed by trials with similar parameter settings (under positive autocorrelation), thus inducing temporal clustering in errors. For example, when the drift rate remains low across several consecutive trials, this is likely to produce a sequence of slow, response-independent errors (e.g., *slow left, slow right, slow left*). A comparable response-independent pattern may arise under low boundaries, although errors will generally be fast (e.g., *fast left, fast right, fast left*). By contrast, when the starting point is biased toward the left boundary, errors tend to be fast and response-specific (e.g., *fast left, fast left, fast left*). Indeed, when simulating data from the autocorrelated DDM, not only do we see the typical inverted-U conditional accuracy function (Figure 1C, right) but critically this model now also predicts temporal clustering of errors (Figure 1D, right).

Going beyond model simulations, in the current work we implemented an effective fitting procedure of the autocorrelated DDM whose parameters fluctuate in an autocorrelated fashion. In contrast to other DDM versions such as the vanilla or across-trial variability DDM, the autocorrelated DDM predicts temporal clustering of errors. In the following, we elaborate on the technical implementation of the autocorrelated DDM, report the recovery of its parameters with as few as 500 trials, and discuss the fit of the three models to 8 empirical datasets, consisting of both human and mouse data.

## Methods

### Accessing the model

To promote the use of the autocorrelated DDM by other researchers interested in measuring fluctuations in DDM parameters we made the model open-source. All code and data is available at https://github.com/robinvloeberghs/Autocorrelated-DDM-and-error-clustering.

### Open science statement

We first conducted an exploratory analysis using the data from Experiment 2B of Desender et al. (2022). We then performed a preregistered confirmatory analysis on seven datasets comprising both human and mouse data. The six human datasets were selected from the Confidence Database (Rahnev et al., 2020), whereas the mouse dataset was obtained from Zang et al. (2026). The pre-registration is available at https://osf.io/x4ngf/registrations. For ease of reading, in the remainder we will discuss the findings from the exploratory analysis together with the seven confirmatory analyses.

### Dataset selection

As pre-registered, we selected datasets from the Confidence Database based on following inclusion criteria: 1) the task design must be compatible with the DDM framework (i.e., speeded 2AFC task with responses given via one of two button presses), 2) each participant completed a minimum of 500 trials (i.e., to allow robust estimation of trial-by-trial fluctuations in the autocorrelated DDM), 3) the main manipulation involved the stimulus strength presented to participants on the screen, and 4) no trial-by-trial feedback was provided. Based on these criteria, a total of six datasets were selected. Combined with the mouse dataset of Zang et al. (2026) and the exploratory dataset of Desender et al. (2022) the full analysis comprised eight datasets.

From the eight studies included, six used Gabor patches as experimental stimuli, one study used the random dot motion task (i.e., participants indicated whether they perceived left-or rightward movement within a cloud of moving dots), and one study asked participants to decide whether the left or right panel contained the most dots. Across studies, the number of subjects varied from 11 to 99, and the number of trials ranged between 500 and 3240. Trials with reaction times shorter than 0.1s or longer than 3s were excluded from the analyses. In total, after excluding these trials the final datasets comprised 223 subjects with 236.567 trials in total. Detailed information about each of the studies can be found in Supplementary Table 1.

### Generative models

#### Vanilla Drift Diffusion Model

The vanilla DDM assumes that on every trial evidence is accumulated until one of two boundaries is reached. How much evidence needs to be accumulated is determined by the boundary separation *a*. The evidence accumulation process itself is assumed to follow a Wiener process:

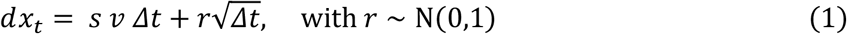

with *dx*_*t*_ representing the change in evidence on each time step *t*. It consists of a deterministic part *s v* Δ*t*, and a random part 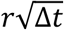. The deterministic part consists of a signed stimulus *s* (between −1 and +1) and the drift rate *v*. The direction and strength of the evidence accumulation directly depends on the signed stimulus *s* and drift rate *v* such that a positive value for *s* on average will steer the evidence accumulation towards to upper boundary, and a negative value to the lower boundary. The random fluctuation is driven by a noise component *r*. The step size in time Δ*t* is 0.001.

On each trial, the evidence accumulation process begins at *z* ⋅ *a*, where *z* is the starting point expressed as a proportion of the boundary separation *a*. Thus, *z* takes a value between 0 and 1. A starting point of *z* = 0.5 indicates no bias, as the process begins exactly halfway between the lower and upper decision boundaries.

This generative mechanism gives rise to both a response and a reaction time on a given trial. The reaction time is determined as the number of time steps needed to reach one of the boundaries with the non-decision time *T*_*er*_ added. The response is determined by whether the upper or lower boundary is reached. Together, the vanilla DDM has four parameters (*v, a, z, T*_*er*_).

#### Across-trial variability Drift Diffusion Model

The across-trial variability DDM extends the standard DDM by assuming that both drift rate and starting point are randomly drawn from a distribution (i.e., *IID* noise). On each trial the drift rate is drawn from a normal distribution centered around the true drift rate *v*:

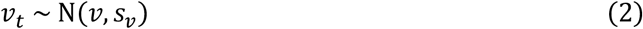

The starting point is drawn from a uniform distribution around true *z* with range *s*_*z*_:

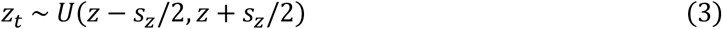

With two additional parameters compared to the standard DDM, the across-trial variability DDM has 6 parameters (*v, a, z, T*_*er*_, *s*_*v*_, *s*_*z*_).

#### Autocorrelated Drift Diffusion Model

Whereas the across-trial variability DDM assumes *IID* noise, the autocorrelated DDM models the trial-to-trial fluctuations of drift rate, starting point, and boundary as a first-order autoregressive model (AR1). The drift rate, starting point, and boundary on a given trial *t* consist of a constant part (*v, z*, and *a*), and a fluctuating part (*γ*_*t*_, *ζ*_*t*_, and *α*_*t*_),

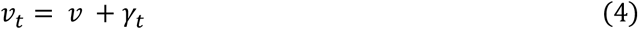

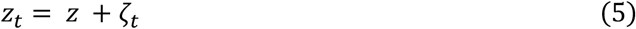

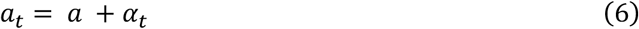

The constant part captures the overall mean of the parameter over all trials, whereas the fluctuating part represent the trial-by-trial fluctuations around this mean. These trial-to-trial fluctuations (*γ*_*t*_, *ζ*_*t*_, and *α*_*t*_) are explicitly estimated and fluctuate according to an AR(1) model from trial *t*-1 to trial *t*:

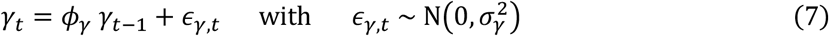

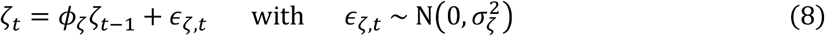

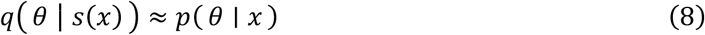

where *ϕ*_*γ*_, *ϕ*_*ζ*_, and *ϕ*_*α*_ are the autoregressive coefficients and 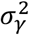, 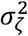, and 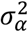 the innovation variances. The autoregressive coefficients govern the strength of the autocorrelation of a time series, whereas the innovation variance acts as a scaling parameter. Note that the intercept is fixed to zero for all three AR(1) models. This implies that the mean of trial-to-trial fluctuations is zero as well. A time series with a generative mean different from zero would result in trade-offs with the constant parts (*v, z*, and *a*), and should therefore be avoided. To ensure that the sum of *z* and *ζ*_*t*_ stays between 0 and 1 a sigmoid transformation is applied. In addition to an estimation of *γ*_*t*_, *ζ*_*t*_ and *α*_*t*_ on each trial *t* = 1, …, *T*, this model has 10 parameters 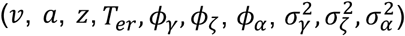.

Hereafter, a distinction between three types of parameters will be made: the *shared* parameters, *hyper*parameters, and *local* parameters. First, the shared parameters consist of the standard DDM parameters: drift rate *v*, starting point *z*, boundary *a*, and non-decision time *T*_*er*_. These parameters remain constant across all trials for a participant and are directly employed in the generative mechanism, explained above, to produce reaction times and responses. The hyperparameters, by contrast, govern the autoregressive process AR(1) underlying the fluctuating part of drift rate *γ*_*t*_, starting point *ζ*_*t*_ and boundary *α*_*t*_. They include the autoregressive coefficients *ϕ*_*γ*_, *ϕ*_*ζ*_, and *ϕ*_*α*_, as well as the error variances 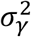, 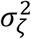, and 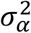. The local parameters are the trial-to-trial fluctuations for drift rate *γ*_1:*T*_, starting point *ζ*_1:*T*_, and boundary *α*_1:*T*_.

### Simulating from Drift Diffusion Models

From these three DDMs we simulated responses and reaction times to assess whether temporal clustering of errors emerges. From each model 50 agents were simulated with 10000 trials per agent. The generative parameters were set to *v* = 1.5, *z* = 0.5, *a* = 1, and *T*_*er*_ = 0.2. Across-trial variability was introduced with *s*_*v*_ = 1.25 for the drift rate and *s*_*z*_ = 0.75 for starting point in the across-trial variability DDM. For the autocorrelated DDM, drift rate, starting point, and boundary parameters followed autoregressive processes with coefficient *ϕ* = 0.99. The corresponding innovation variances were 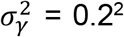, 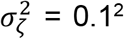, and 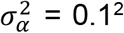. All parameter values were chosen to yield datasets with an average accuracy of approximately 0.75. The results of these simulations are shown in Figure 1.

### A signature for clustered errors

To investigate the temporal clustering of errors in both empirical participants and simulated agents, we constructed a repetition matrix using a subset of trials in which two consecutive responses were errors. Category pairs were created by crossing reaction time (fast vs. slow, defined as below or above the median) with response (left vs. right). The frequencies of the category pairs were then normalized into conditional probabilities such that each column summed to 1. Under this normalization, a value of 0.25 corresponds to a chance-level baseline, while accounting for differences in the marginal frequencies of the different categories. This procedure was applied separately to each agent, and the final repetition matrix shows the average conditional probabilities across all agents. The presence of fast and slow errors was investigated through the conditional accuracy function, where accuracy was computed as a function of reaction time quantiles calculated per agent.

### Estimation of Drift Diffusion Model parameters

To estimate the DDM parameters we relied on Amortized Bayesian Inference (ABI), which is implemented in the Python package Bayesflow (version 1.0) (Radev et al., 2023). ABI enables the estimation of parameters for models with high complexity, using neural networks. The key tenet behind ABI is to simulate many datasets from the generative model and train a neural network to approximate the posterior distribution over parameters given observed data. The workflow for ABI is illustrated in Figure 2. During the training phase, a prior over the parameters *θ* ~ *p*(*θ*) is specified together with a observation model *X* ~ *p*(*X*|*θ*). From this observation model, conditional on the sampled parameters, data is drawn (responses and reaction times). Repeating this procedure yields a large number N of simulated datasets, which together with the underlying generative parameters 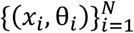, are passed to the inference network. Prior to entering the inference network, the datasets are processed by a summary network. The summary network maps each dataset *x*_*i*_ to a fixed-size vector of approximately sufficient summary statistics *s*(*x*_*i*_), where *s*(⋅) denotes the transformation performed by the network. This ensures that the inference network receives datasets all with consistent input dimensions. The inference network then learns the data-posterior mapping by minimizing the Kullback-Leibler (KL) distance between the true and approximate posterior distribution (Kullback & Leibler, 1951; Radev et al., 2022):

**Figure 2:**
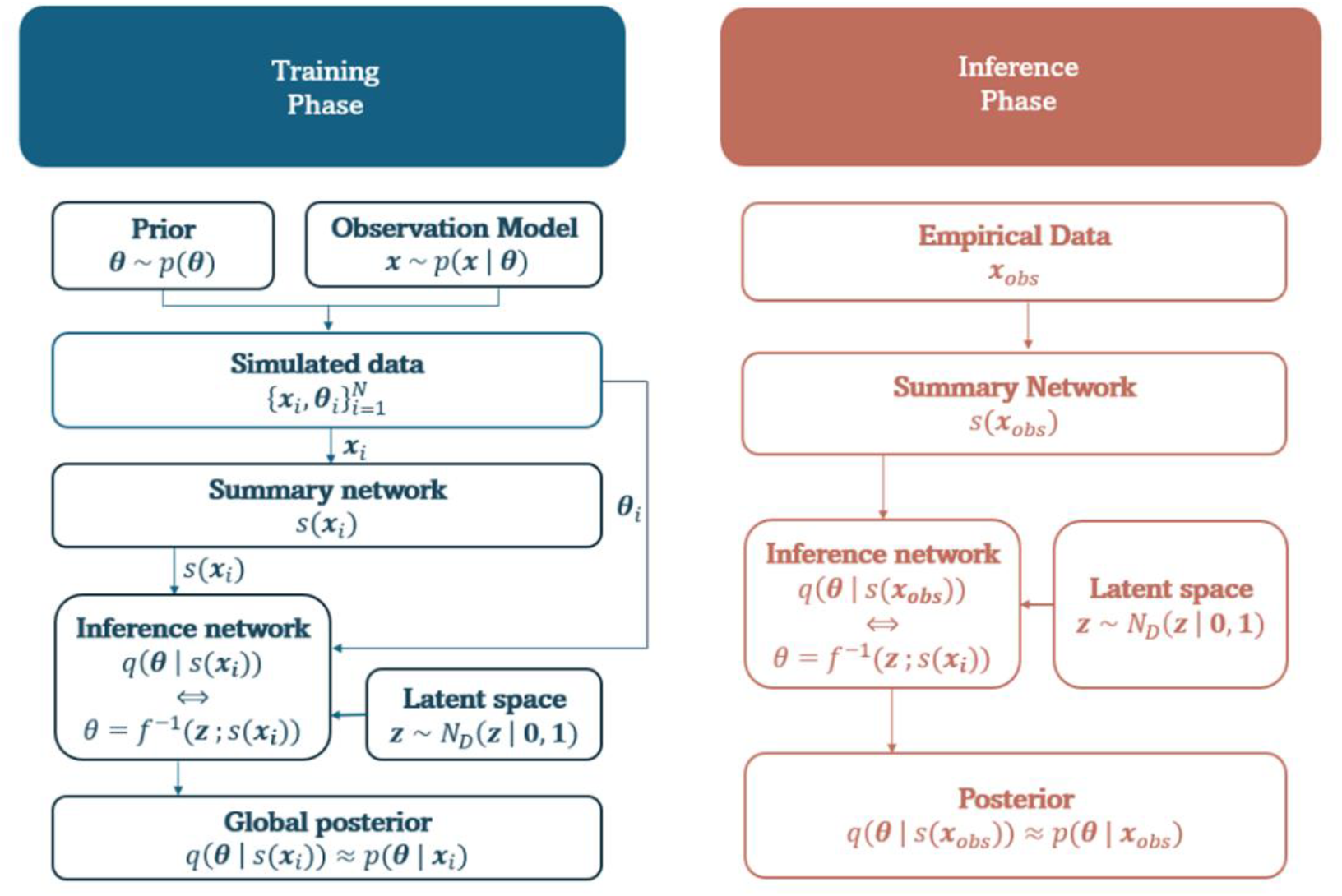
Workflow overview of Amortized Bayesian Inference. During the training phase, a set of neural networks learns the relationship between observed data *x* and the underlying generative parameters θ to estimate the posterior distributions. In the inference phase, new observed data *x*_*obs*_ is passed through the trained model to produce the posterior estimates almost instantly. Figure adapted from Wu et al. (2026), with permission.

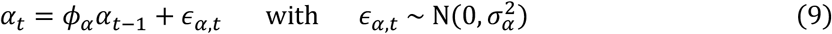

Sampling from this complex, high-dimensional posterior distribution *q*( *θ* ∣ *s*(*x*)) can be challenging. To address this challenge, ABI employs normalizing flows (Kobyzev et al., 2021), which learns an invertible transformation between a simple distribution (e.g., a normal Gaussian) and the complex target posterior. By learning this transformation, the model can sample from the base distribution and map the samples through the invertible function to obtain approximate posterior samples:

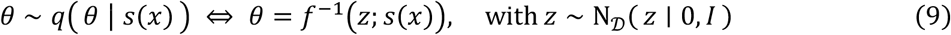

After an extensive training phase, the inference process for new datasets becomes highly efficient, with almost instantaneous posterior sampling. The empirical data *x*_*obs*_ is passed forward through the trained network, producing the approximate posterior distribution *q*( *θ* ∣ *s*(*x*)) from which the posterior samples can be drawn. This efficiency is one of the key advantages of ABI: the computational cost is concentrated in the training phase, allowing for very fast inference across multiple participants or experimental conditions.

For the training phase, wide uniform prior distributions were chosen that cover the whole range of empirically plausible values (Suppl. Table 2). The drawn parameter values are then used to generate responses and reaction times for each of the three generative models described above. The reaction times are log transformed for model training as this improves convergence (Wu et al., 2026). The number of trials per simulated subject ranged from 350 to 3250, matching the trial counts in the empirical datasets.

The next step involves defining the neural network architecture. This consists of three main components: a hierarchical summary network, a local inference network for the local parameters, and a global inference network, for both the shared and hyperparameters.

The summary network consists of a two-level LSTM-based architecture. The first level of the summary network comprises three LSTM (Long short-term memory) layers with decreasing hidden dimensions (512, 256, 128), each returning sequences, which allows for the preservation of temporal information across input trials. The second level further compresses the representation via a single LSTM layer with 64 hidden units, which outputs a fixed-length vector. The summary network for the vanilla and across-trial variability DDM consisted of a permutation-invariant Set Transformer architecture (Lee et al., 2019)

The inference component is split for local and global posterior approximations. The local inference network targets trial-by-trial fluctuations in drift rate, and starting point, and boundary, and is implemented using an invertible neural network with three output parameters and eight interleaved coupling layers. The global inference network, in contrast, captures both hyperparameters and shared parameters. It outputs ten parameters in total—six corresponding to hyperparameters and four to shared parameters—using twelve interleaved coupling layers.

These components are integrated via a two-level amortized inference architecture, in which the local and global networks are conditioned on the hierarchical summary representations extracted from the data.

All DDMs were trained for 500 epochs, each consisting of 1000 iterations, using a batch size of 32. On GPU-accelerated hardware (NVIDIA V100 GPU, 4 CPU cores), training times were approximately 8 hours for the vanilla DDM, 9 hours for the across-trial variability DDM, and 120 hours for the autocorrelated DDM.

In the empirical data, after training the three DDM variants, each model was fitted to every participant separately. The posterior mean for each parameter was calculated based on 50 posterior samples.

### Parameter recovery and model checks

The parameter recovery for each DDM variant was performed on 1000 simulated agents, with 500 trials per agent, and the generative parameters drawn from the same uniform distributions as in the training procedure (see Suppl. Table 1). After estimation, the posterior mean for each parameters was calculated on 50 posterior samples. To assess convergence of the model the losses over iterations were visually inspected.

Simulation-Based Calibration (SBC) was used to investigate whether the inference method is well-calibrated (i.e., no biases are present in the posteriors) (Säilynoja et al., 2022; Talts et al., 2020). To implement SBC, parameters are first sampled from the prior and used to generate datasets. Next, the posterior distributions are estimated for each simulated dataset. From these posteriors, samples are drawn and the rank of the *true* parameter within these samples is computed. If the inference method is well-calibrated, the ranks aggregated over many simulations, should follow a uniform distribution. Deviations from uniformity indicate miscalibration, such as bias or posteriors being too wide or narrow. For reliable estimation of confidence intervals in the rank histograms, it is recommended that the ratio of simulated agents to posterior samples be at least 20, which is satisfied in this case.

### Model fit

Model fit was evaluated by comparing predicted performance to determine which model best accounted for the empirical data. After fitting each DDM variant to the empirical data, we simulated 50 new datasets for each subject using parameters randomly drawn from the estimated posterior distribution. Based on the sampled hyperparameters, new trial-to-trial fluctuations for drift rate, starting point, and boundary were generated. This procedure enables to propagate posterior uncertainty into the predictive simulations (i.e., posterior predictive checks; Gelman et al., 1996). To ensure sufficiently stable predictions, the empirical stimulus difficulty sequence was repeated 20 times and randomly shuffled prior to simulation. For each simulated dataset, we computed the conditional accuracy function (CAF) and the repetition matrix. For each participant, these measures were then averaged across the 50 simulated datasets. Model performance was quantified by calculating, for each participant, the correlation between the averaged simulated (i.e., derived from data generated under estimated parameters) and empirical CAF and repetition matrix.

To test whether the correlations between simulated and empirical CAFs and repetition matrices differed across DDM variants, we fitted linear mixed models with DDM variant as a predictor of the correlations. Pairwise differences between DDM variants were assessed using two-sided post hoc contrasts with a Tukey-Kramer correction for multiple comparisons. First, we analyzed all datasets *combined*, while accounting for the nesting of subjects within datasets by using nested random effects. In a second analysis, we applied a linear mixed model to each dataset *individually*.

Note that traditional information criteria such as AIC and BIC cannot be computed within the Amortized Bayesian Inference (ABI) framework because it does not provide a closed-form likelihood function. Although Bayesian model comparison (BMC) using neural networks is in principle possible, this approach is not recommended. BMC is known to be highly fragile, particularly when applied to neural networks (Elsemüller et al., 2024).

## Results

### Parameter recovery

To assess parameter recovery, we simulated 1000 agents performing 500 trials for each model variant. For each agent, posterior means were computed from 50 posterior draws per parameter. Recovery was quantified as the correlation between the true parameter values and the inferred posterior means.

In the vanilla DDM, the shared parameters – drift rate, starting point, boundary separation, and non-decision time – show perfect recovery (all *r* > 0.99) (Suppl. Figure 1A). Also in the across-trial variability DDM these parameters show a near-perfect correlation between the true and inferred posterior means (all *r* > 0.98). The across-trial variability parameters, which are known to be difficult to estimate reliably, also show reasonable recovery overall, although recovery is markedly better for drift rate variability *s*_*v*_ (with a correlation *r* = 0.88) than for starting point variability *s*_*z*_ (*r* = 0.44).

In the autocorrelated DDM, the shared parameters can be accurately recovered, with correlations exceeding .94 for all parameters (Figure 3A). For the hyperparameters, correlations are at least 0.82 for the autoregressive coefficients *ϕ* and 0.94 for the innovation standard deviations *σ* (Figure 3B). The recovery of the local parameters (i.e., the trial-to-trial fluctuations in drift rate, starting point, and boundary) is shown in Figure 3C. The correlation between the generative and inferred trajectories depends on generative parameters of the time series. More specifically, for all three parameters the recoverability is higher when the generative autoregressive coefficient *ϕ* is large. When *ϕ* is small, the time series contains less systematic autocorrelation and instead resembles random noise, making recovery more difficult. This can partly be countered when a low value for *ϕ* is combined with a high innovation standard deviation σ. Since *σ* acts as a scaling parameter, larger values produce more pronounced fluctuations in the time series, which in turn makes it easier for the model to detect and recover the underlying trajectory.

**Figure 3:**
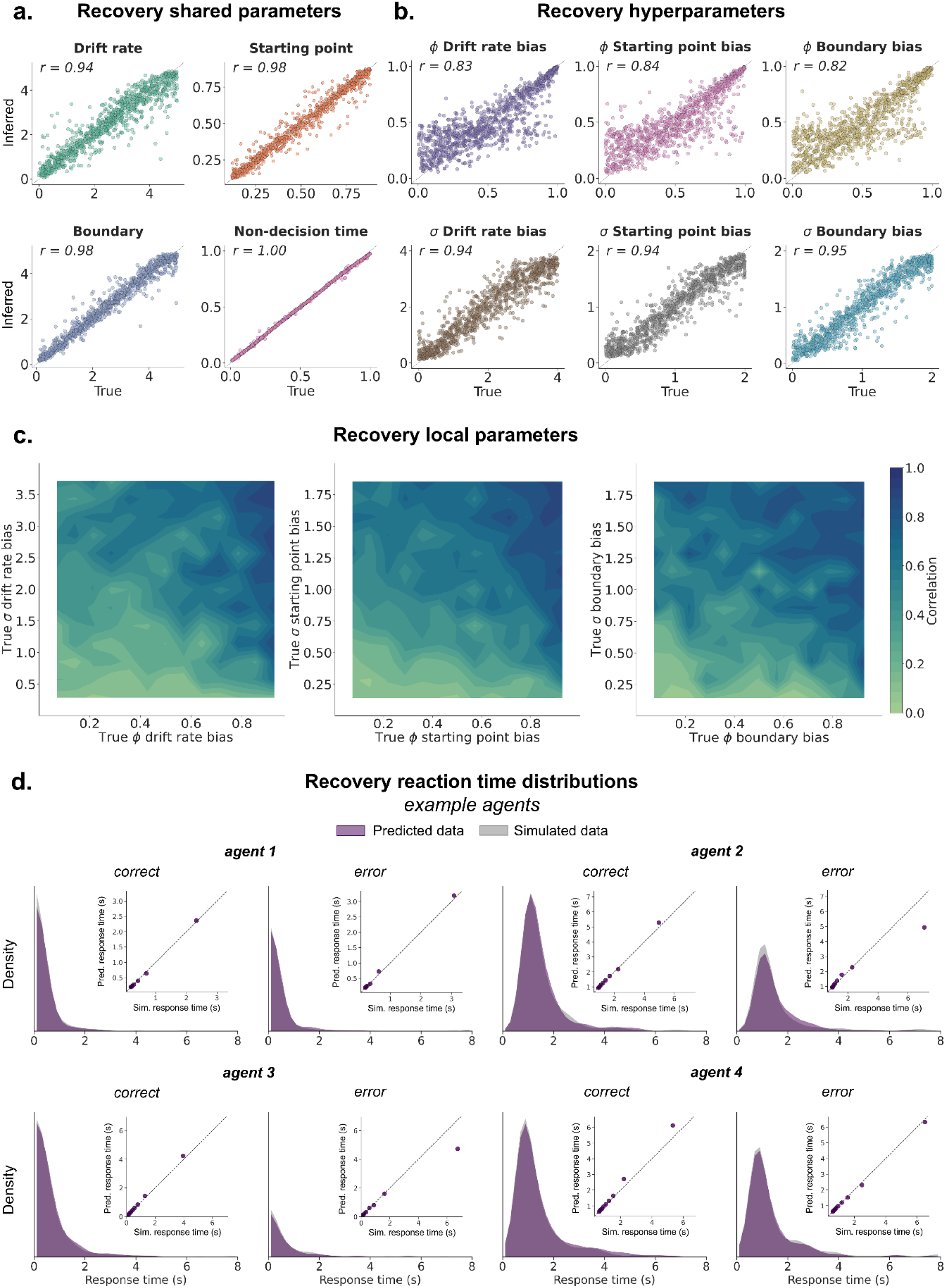
Overview of the parameter recovery for the autocorrelated DDM using Amortized Bayesian Inference. **a-b**. Parameter recovery of the posterior means for the shared parameters (a) and hyperparameters (b). **c**. Recovery of the trial-to-trial fluctuations (i.e., local parameters), quantified as the correlation between the true values and the inferred posterior means, is shown as a function of the generative AR(1) parameters. For all three parameters we see an excellent recovery with a high correlation between the true and estimated fluctuations when the true *ϕ* and/or *σ* is high. A time series is generated with a low autoregressive coefficient and innovation variance will resemble random noise, which makes it hard for the model to accurately recover. **d**. Predicted (i.e., based on estimated parameters) and simulated (i.e., based on generative parameters) and reaction time distributions for correct and error trials from four example agents. The inset shows quantile-quantile plots comparing the predicted and simulated reaction time distributions across 10 quantiles (dots). The identity line indicates perfect correspondence between the predicted and simulated reaction time distributions.

The autocorrelated DDM shows a great correspondence between the true and inferred reaction time distributions, both for correct and error trials (Figure 3D). Lastly, using Simulation-Based Calibration (SBC), we confirmed that no strong biases are present in the posterior distributions for most parameters, as indicated by the approximately uniform rank statistics histograms (Suppl. Figure 2). Only for non-decision time *T*_*er*_ we do see a violation, with distinct peaks at the extremes, suggesting that the non-decision time is frequently under- or overestimated. However, as the posterior mean for *T*_*er*_ showed a near-perfect recovery (*r* ~ 1, Figure 3A), this most likely is the consequence of an extremely narrow posterior for non-decision time (average posterior standard deviation = 0.004) compared to the other parameters (average posterior standard deviation > 0.196), with very small deviations from the true posterior mean leading to extreme ranks.

### Fitting empirical data

Given the main focus of the current work is the development of the autocorrelated DDM, we here first focus on its fit to empirical data and its estimated parameters. For both the shared parameters and the hyperparameter we see substantial individual differences in the example dataset of Desender et al. (2022) (Figure 4A-B). Most interestingly, whereas starting point and boundary show highly autocorrelated fluctuations over trials (median *ϕ* = 0.91 for starting point, median *ϕ* = 0.92 for boundary), the fluctuations for drift rate are much less autocorrelated (median *ϕ* = 0.48). Indeed, visual inspection of two example participants suggests that drift rate fluctuations appear to behave more as random noise compared to the smooth autocorrelated timeseries for starting point and boundary (Figure 4C). The empirical fits for the vanilla DDM and across-trial variability DDM are shown in Supplementary Figure 3. Lastly, for the same two example participants, reaction time distributions simulated using the estimated parameters closely reproduced the empirical distributions, demonstrating that the autocorrelated DDM provided a good fit to the observed data (Figure 4D).

**Figure 4:**
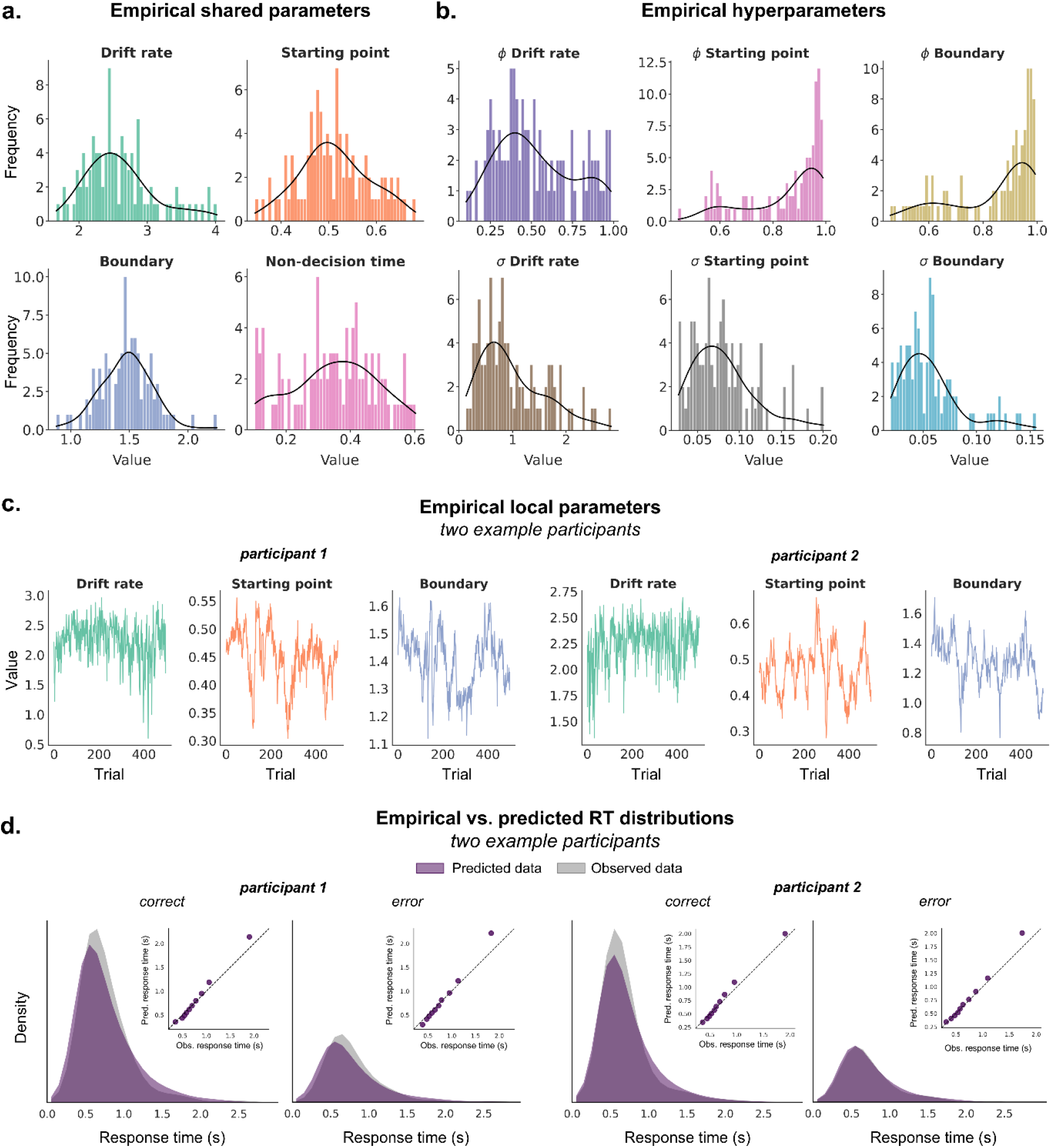
Fitting the autocorrelated DDM to the example dataset of Desender et al. (2022) a-b. Empirical fits of 99 participants for the shared (a) and hyperparameters (b). **c**. The estimated trajectories for drift rate, starting point, and boundary for two example participants. The trajectories show the constant (*v, z*, and *a*) and fluctuating part (*γ*_*t*_, *ζ*_*t*_, and *α*_*t*_) of each parameter together. Note that the evidence accumulation on each trial is defined as the product of the stimulus value (rescaled between –1 and 1) and a slope parameter. The fluctuating drift rate values shown here correspond to the slope parameter. **d**. Predicted (i.e., based on estimated parameters) and observed reaction time distributions for correct and error trials from the same two example participants as in Figure 4C. The inset shows quantile-quantile plots comparing the predicted and observed reaction time distributions across 10 quantiles (dots). The identity line indicates perfect correspondence between the predicted and simulated reaction time distributions.

### Temporal clustering of errors

Having fitted the three different versions of the DDM to empirical data, we next investigated the main hypothesis: can the autocorrelated DDM capture temporal clustering of errors in empirical data? The repetition matrix averaged over all empirical subjects of the 8 datasets clearly shows that, indeed, errors do not occur randomly in time (Figure 5A). Rather, there is a clustering of fast response-specific errors (e.g., fast left – fast left) in the upper left quadrant of the repetition matrix. This presumably follows from autocorrelated fluctuations in starting point, as simulated data with only the starting point fluctuating according to an AR(1) process exhibit the same pattern (Suppl. Figure 4A). In addition to the fast response-specific errors, we also see a clustering of slow response-independent errors (e.g., slow left – slow right). In simulated data this type of clustering occurs following AR(1) fluctuations in drift rate, boundary, or both (Suppl. Figure 4B-D).

**Figure 5:**
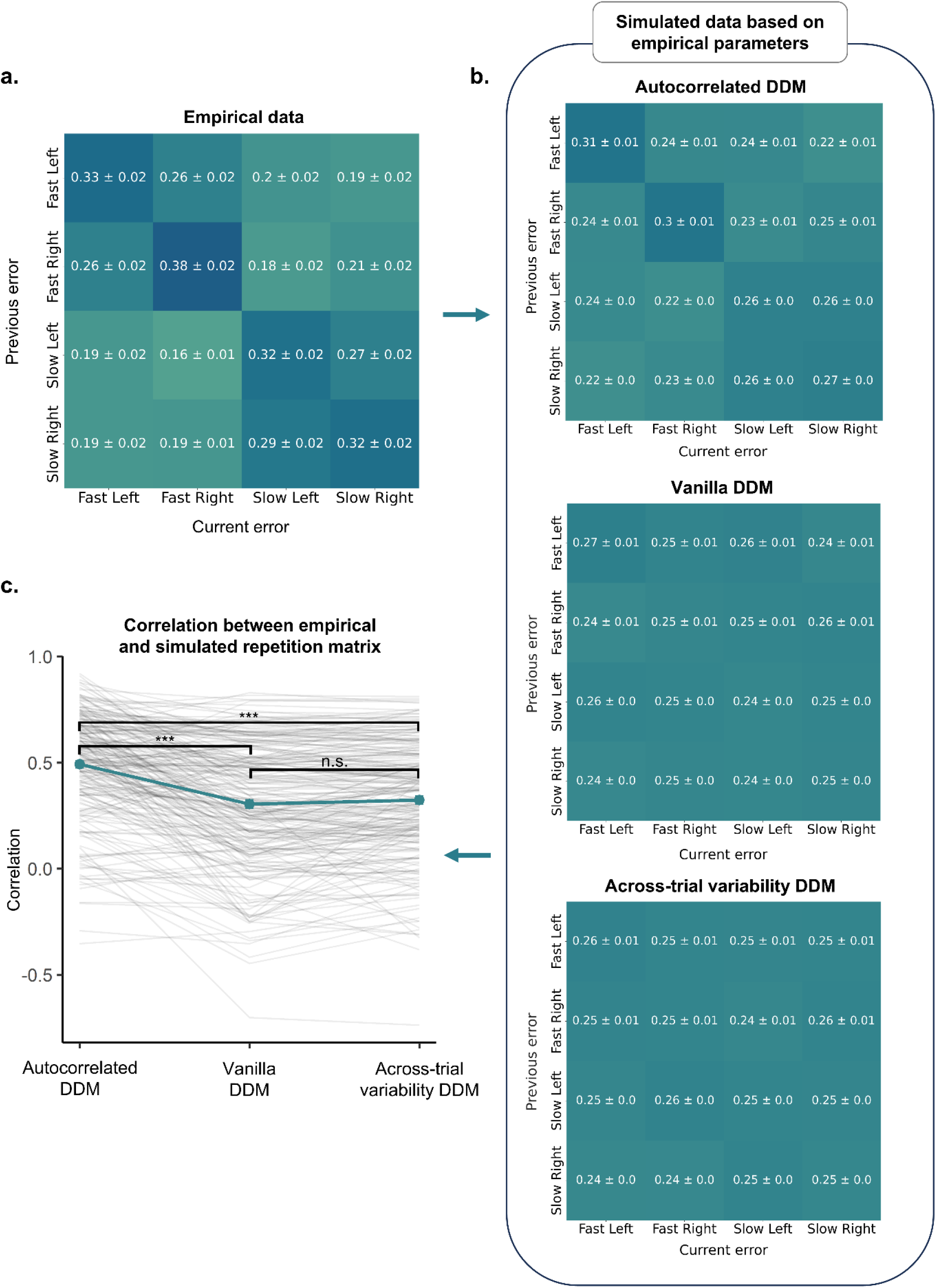
Temporal clustering of errors across all eight empirical datasets is best explained by the autocorrelated DDM. **a**. The repetition matrix shows the proportion of current–previous error co-occurrences, with 0.25 representing chance level. For the empirical data, the repetition matrix was computed separately for each participant and then averaged across all 223 participants. The data (± standard error) show clear clustering for fast response-specific errors (e.g., fast left – fast left), and slow response-independent errors (e.g., slow left – slow right). **b**. The repetition matrices for the three DDM variants after simulating data with parameter values drawn from the posterior distributions. Albeit a smaller effect, the repetition matrix for the autocorrelated DDM exhibits the same clustering pattern as observed in the empirical data. **c**. For each participant, we tested how well the empirical repetition matrix corresponded with each of the three simulated repetition matrices using correlations. The correspondence between empirical data and data simulated under the autocorrelated DDM are significantly higher compared to the other two (autocorrelated vs. vanilla: *p* < .001, autocorrelated vs. across-trial variability: *p* < .001, across-trial variability vs. vanilla: *p* = .331), indicating that the error clustering in empirical data is best explained by the autocorrelated DDM. The average correlation over all participants with error bars (± standard error) is shown. The lines represent the data of individual participants. At first glance, it may seem surprising that the correlations for the vanilla DDM and across-trial variability DDM are not centered around zero, given that neither model can, in principle, account for clustering (i.e., repetition matrices should be around .25). However, participants often have a biased starting point (z ≠ 0.5), which induces a specific structure in the repetition matrix (Suppl. Figure 4E). When we simulate data using these empirical parameter values, the same structure emerges in the simulated repetition matrix, leading to a positive correlation. Note: n.s. = *p* > .05, *** = *p* < .001.

Using the empirical parameter estimates obtained from fitting the three DDM variants, we simulated 50 new datasets for each subject by randomly sampling parameter values from the posterior distribution for each simulation. For each simulated dataset, we computed the repetition matrix and then averaged these matrices across the 50 simulations. Finally, we correlated the resulting average repetition matrix with the corresponding empirical repetition matrix for each subject (Figure 5B–C).

First, when analyzing these correlations *across* all eight datasets in a linear mixed model with nested random effects, we found that the correspondence between empirically observed and model-predicted clustering of errors was significantly different between the three DDM variants 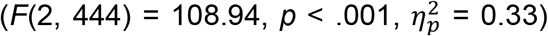 (Figure 5C). Post-hoc comparisons show that the autocorrelated DDM outperformed the other models, with significantly better correspondence compared to the vanilla DDM (*β* = 0.193, SE = 0.014, *t*(444) = 13.435, *p* < .001, *d* = 1.272, CI(95) = [0.159, 0.227]), and compared to the across-trial variability DDM (*β* = 0.172, SE = 0.014, *t*(444) = 12.013, *p* < .001, *d* = 1.138, CI(95) = [0.139, 0.206]). As expected, there was no significant difference between the across-trial variability DDM and the vanilla DDM (*β* = 0.02, SE = 0.014, *t*(444) = 1.421, *p* = .331, *d* = 0.135, CI(95) = [-0.013, 0.054]). Note that repeating the analysis using Fisher z-transformed correlations as the dependent variable resulted in nearly identical results. Out of 223 participants, 164 participants (73.54%) showed the highest correspondence for the autocorrelated DDM, whereas the vanilla DDM and across-trial variability DDM were the preferred model in only 22 and 37 participants (9.87% and 16.59%), respectively. The correspondence between empirical and fitted conditional accuracy function for each model can be found in Supplementary Figure 5.

In the previous section, we have shown that *across* all eight datasets the autocorrelated DDM provided the best fit to the empirical repetition matrices. Next, we investigated the correspondence between the simulated and empirical repetition matrices for the three DDM variants for each dataset *individually* (Figure 6). Using linear mixed models, we found that in every dataset the correlation was significantly higher for the autocorrelated DDM than for both the vanilla DDM and the across-trial variability DDM. In contrast, no significant differences were observed between the vanilla DDM and the across-trial variability DDM in any of the eight datasets. The simulated and empirical repetition matrices for each dataset separately are shown in Supplementary Figure 6. Together, these findings indicate that the autocorrelated DDM most accurately captures the temporal clustering of errors observed in the empirical data, both when all eight datasets are analyzed jointly and when they are examined individually.

**Figure 6:**
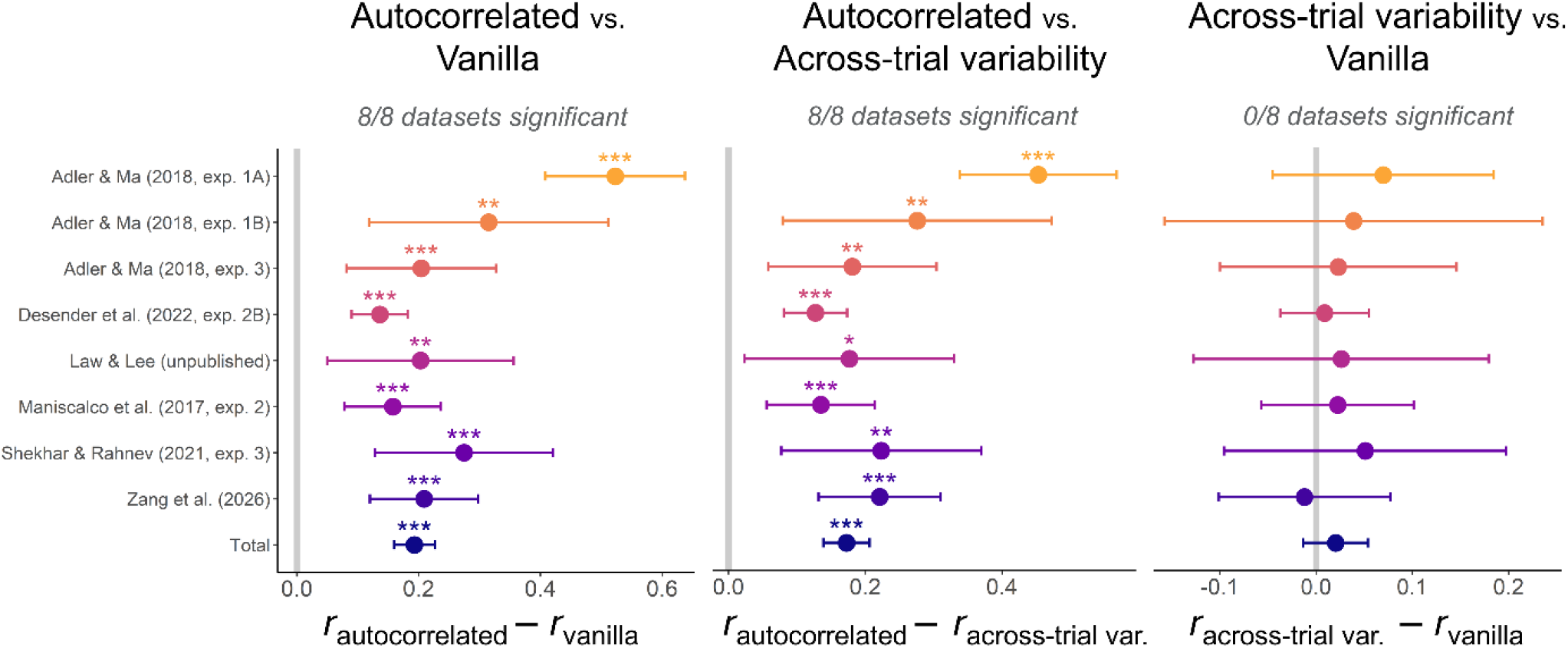
The autocorrelated DDM provides the best fit for the temporal clustering of errors in all eight datasets, consisting of both human and mouse data. Linear mixed models comparing the correlation between empirical and simulated (i.e., generated based on estimated DDM parameters) repetition matrices for the three DDM variants showed that, in every dataset, the autocorrelated DDM produced significantly higher correlations than both the vanilla DDM and the across-trial variability DDM. No significant differences were observed between the vanilla DDM and the across-trial variability DDM. The figure shows the parameter estimates for the post hoc contrasts together with their 95% confidence intervals. Estimates were obtained from models fitted separately to each dataset. The estimate at the bottom, labeled “Total”, is based on a model fitted to all datasets simultaneously while accounting for the nesting of subjects within experiments. Note: * = *p* <.05, ** = *p* <.01, *** = *p* <.001.

## Discussion

One of the most widely used computational models of decision-making is the drift diffusion model (DDM). In its simplest four-parameter formulation, the vanilla DDM accounts for key behavioral phenomena, including difficulty effects and speed-accuracy trade-offs. Extensions of the model introduce across-trial variability by adding random noise to the drift rate and starting point, enabling the model to explain fast and slow errors that the vanilla DDM cannot capture. This variability is typically implemented as *independent and identically distributed* (IID) draws, for example, normally distributed noise for drift rate and uniformly distributed noise for starting point. A direct consequence of this *IID* assumption is that errors occur randomly in time. Intuitively, if the starting point on a given trial is very close to the incorrect boundary, an error is likely; however, because trial-to-trial variability is independent, the starting point on the subsequent trial may be far away from that boundary, reducing the probability of another error. In contrast, empirical data commonly exhibits temporal clustering of errors, with errors tending to occur in sequences rather than in isolation (Dutilh, Vandekerckhove, et al., 2012; Gilden & Wilson, 1995; Malykke et al., 2026). To resolve this mismatch between model assumptions and observed behavior, we here propose an autocorrelated DDM in which drift rate, starting point, and boundary fluctuate across trials according to a first-order autoregressive (AR1) process. First, in simulations we demonstrate that the autocorrelated DDM is able to explain temporal clustering of errors, whereas both the vanilla DDM and the across-trial variability DDM are unable to generate these patterns. Consistent with these simulation results, analyses of eight empirical datasets, comprising both human and mouse data, show that the autocorrelated DDM exhibits the closest correspondence to the empirical repetition matrices. Furthermore, when each dataset is analyzed individually, the autocorrelated DDM provides the best account of the temporal clustering of errors across all eight datasets.

Different from the classical implementation of the DDM, which assumes potentially noisy but otherwise stable parameters across time, the parameters in the autocorrelated DDM can fluctuate over time. This approach aligns with a broader shift in cognitive neuroscience toward modeling *time-varying* internal states (Ashwood et al., 2022; Cowley et al., 2020; Miletic et al., 2025; Schumacher et al., 2023; Urai, 2025; Vloeberghs et al., 2025, 2026). As reviewed by Urai (2025), what has often been described as unexplained variance in previous modeling efforts, can often be captured in such time-varying models as genuine signal driving variability in behavior. Here, we specifically assume that parameters vary according to an autoregressive process. Note that while fluctuations in parameters may potentially be driven by multiple sources (e.g., attention, arousal, mood), the model itself remains agnostic regarding what drives these fluctuations. To examine whether these fluctuations are systematically driven by external factors—such as for instance boundary shifts following an error response—the framework can be extended in a straightforward manner to incorporate additional covariates, such as previous trial accuracy to investigate post-error slowing (Dutilh, Vandekerckhove, et al., 2012).

The current work is not the first to adapt the DDM such that it can account for temporal changes in model parameters. Whereas some older work provided very coarse estimates by estimating DDM parameters separately for blocks of trials (Dutilh et al., 2009; Evans & Brown, 2017; Vermeylen et al., 2023), more recent work estimated categorically distinct latent states, with each state characterized by a distinct set of parameters (Gunawan et al., 2022), or constrained how parameters evolve over time by specifying pre-defined (exponential or power) functions (Alister & Evans, 2026). This line of work is highly relevant since not accounting for fluctuations in computational parameters can lead to biased parameter estimates (Alister & Evans, 2026; Vloeberghs et al., 2025). Moreover, ignoring parameter fluctuations can lead to systematic misinterpretations of behavioral effects. For example, consider a decision boundary that gradually decreases over the course of the experiment, as in the example (empirical) participant in Figure 4C. Early in the experiment, a higher boundary produces slower but more accurate responses, whereas later a lower boundary yields faster but less accurate responses. Because post-error adjustments are typically quantified as the difference in response times between post-error and post-correct trials, such global downward trend in the boundary can spuriously manifest as post-error speeding. Crucially, this apparent effect does not reflect a genuine trial-by-trial adjustment following errors, but rather a confound introduced by slow drifts in computational parameters (Dutilh, van Ravenzwaaij, et al., 2012). Confounds such as these can be accounted for within the autocorrelated model by explicitly estimating fluctuations in model parameters.

Our findings have important implications for the post error slowing (PES) literature, where temporal clustering of errors (i.e., what happens after an error response) plays a central role. Traditionally, PES is explained by an adaptive increase in decision boundary following an error to avoid future mistakes (Dutilh, Vandekerckhove, et al., 2012). However, while this account predicts increased accuracy on post-error trials, this effect on accuracy is not always found in empirical data (Hajcak et al., 2003; Hajcak & Simons, 2008). To address this discrepancy, the orienting account proposes that errors trigger a reduced drift rate, leading to post-error slowing and post-error accuracy decrease (Notebaert et al., 2009). Importantly, both accounts assume that the cognitive system *actively* adjusts a computational parameter in response of having made an error. However, our simulations and empirical results suggest that temporal clustering of errors, or a post-error accuracy decrease, can be explained by *randomly* fluctuating parameters (i.e., not actively driven by a covariate such as accuracy on the previous trial). This alternative explanation could be tested by extending the autocorrelated DDM to include previous-trial accuracy as a covariate predicting trial-to-trial parameter changes. Demonstrating that this covariate does not predict parameter changes would provide direct evidence against the dominant accounts of PES.

Being able to estimate trial-to-trial estimates of DDM parameters has clear benefits for empirical work. However, the estimation of the full latent trajectories of the drift rate, starting point, and boundary across all trials does substantially increase model complexity. As such, it becomes less feasible to estimate it with classical estimation techniques such as Markov Chain Monte Carlo (MCMC) or Gibbs sampling. To resolve this issue, we employed Amortized Bayesian Inference (ABI). ABI is a simulation-based inference framework that leverages neural networks to approximate posterior distributions efficiently. Rather than estimating parameters for each dataset individually (as in classical Bayesian methods), ABI learns a global mapping from data to posterior distributions by training on simulated data. Once trained, this mapping can be applied to new datasets at negligible computational cost, enabling near-instant posterior inference. For example, using ABI, Von Krause et al. (2022) applied a DDM to a dataset consisting of 1.2 million participants, which would be unfeasible with the current gold-standard MCMC methods. Using this estimation technique, in the current work we showed that the parameters of the autocorrelated DDM can reliably be recovered, allowing to make inferences about key parameters at single-trial level, even with only 500 trials per agent.

In sum, in the current work we propose the autocorrelated DDM where drift rate, starting point, and boundary fluctuate from trial-to-trial according to an AR(1) process. We show that in contrast to the vanilla DDM and the across-trial variability DDM, the autocorrelated DDM is able to successfully capture the temporal clustering of errors, as observed in eight empirical datasets consisting of both human and mouse data.

