## Supplementary Figures 1-6, Supplementary Tables 1-2 for "Explaining temporally clustered errors with an autocorrelated Drift Diffusion Model"

### Supplementary Materials

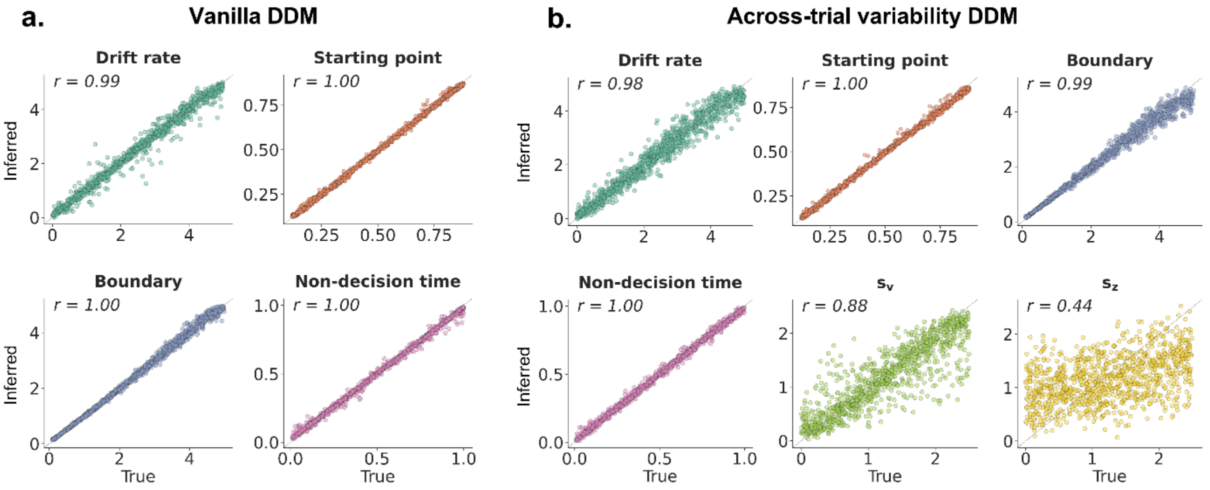


**Supplementary Figure 1: Parameter recovery for the vanilla DDM and across-trial variability DDM.** **a.** The parameter recovery for the vanilla DDM looks excellent with for all parameters an almost perfect recovery (all *r* > 0.99). **b.** Also the across-trial variability for drift rate $s_v$ is highly recoverable, albeit with slightly increased variance around the diagonal (*r* = 0.88). In contrast, the across-trial variability in starting point $s_z$ was more difficult to recover (*r* = 0.44).

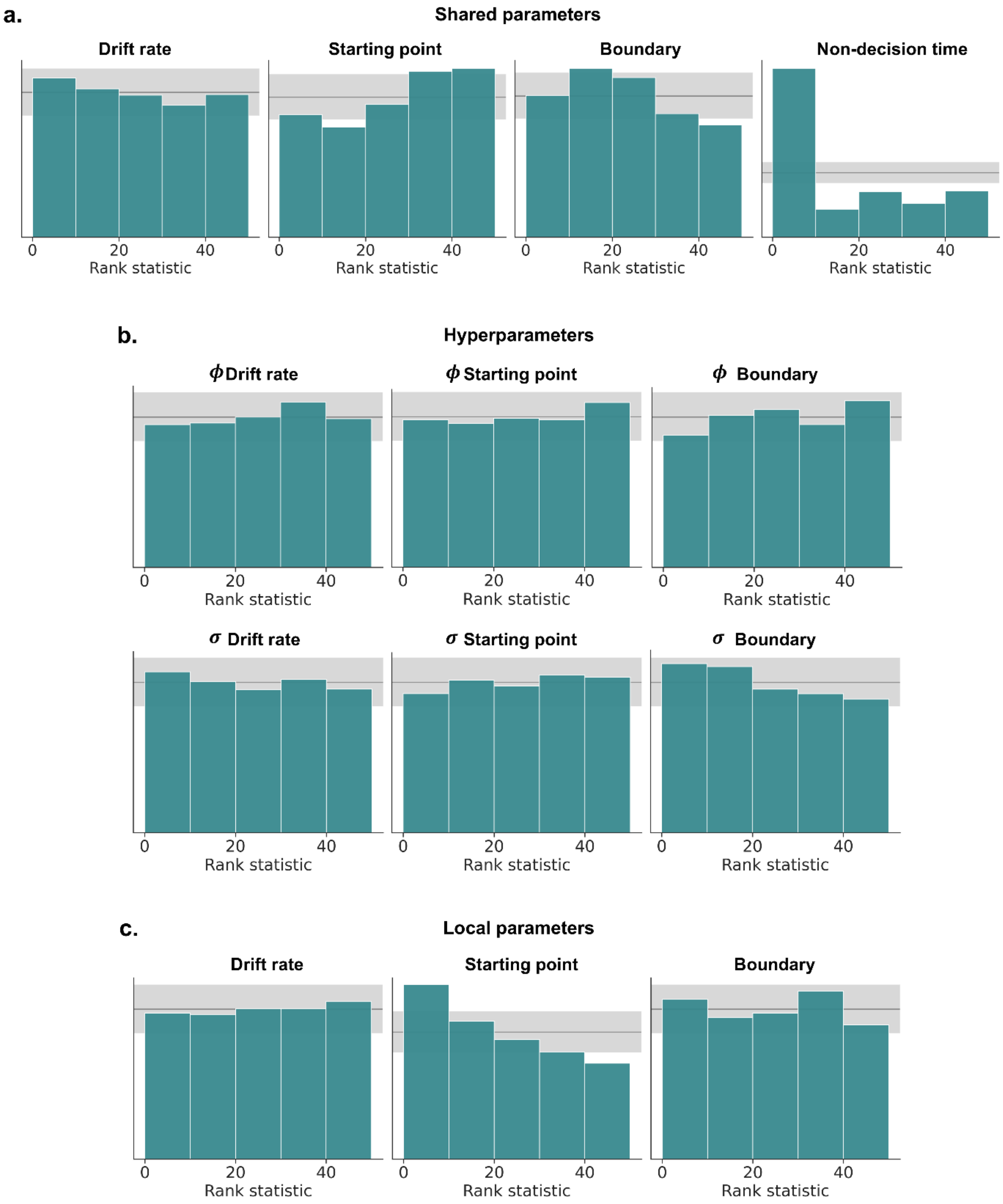


**Supplementary Figure 2: Investigating biased posterior inference using Simulation-Based Calibration (SBC). a.** For the shared parameters, all parameters follow a uniform distribution, except for the non-decision time (gray band indicates confidence intervals). This deviation indicates biased posterior inference. However, model recovery demonstrated perfect recovery for this parameter, suggesting that the deviation is likely due to very narrow posterior distributions. With narrow posterior distributions, even the slightest shift away from the true parameter value can produce extreme rank statistics. **b-c.** The posterior inference is well-calibrated for the hyperparameters (a) and local parameters (c). For the local parameters, the rank statistic histograms are shown for one trial.

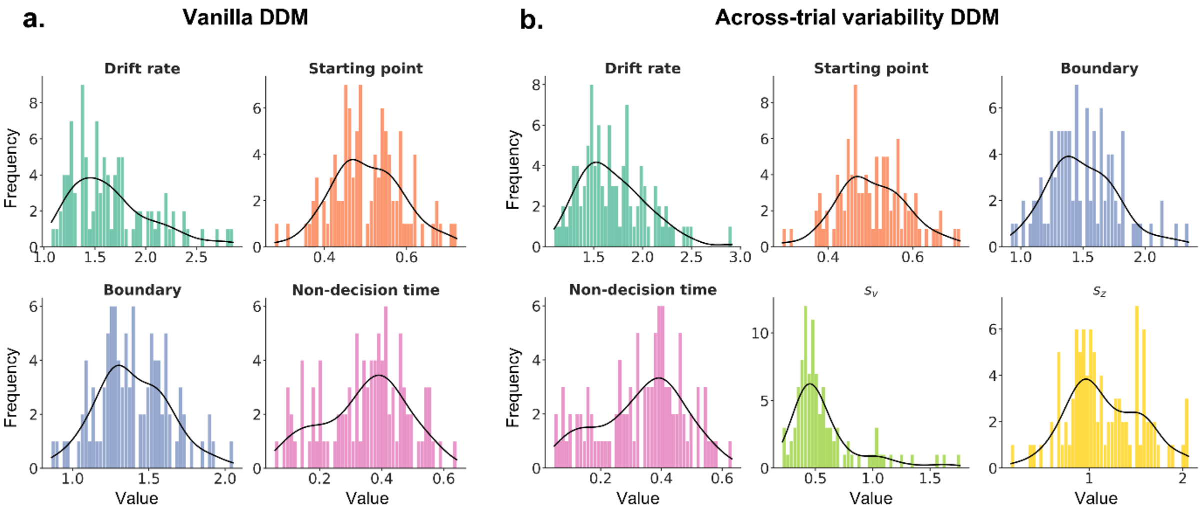


**Supplementary Figure 3: The DDM parameters fitted on the empirical dataset.** The parameters for the vanilla DDM (a) and the across-trial variability DDM (b).

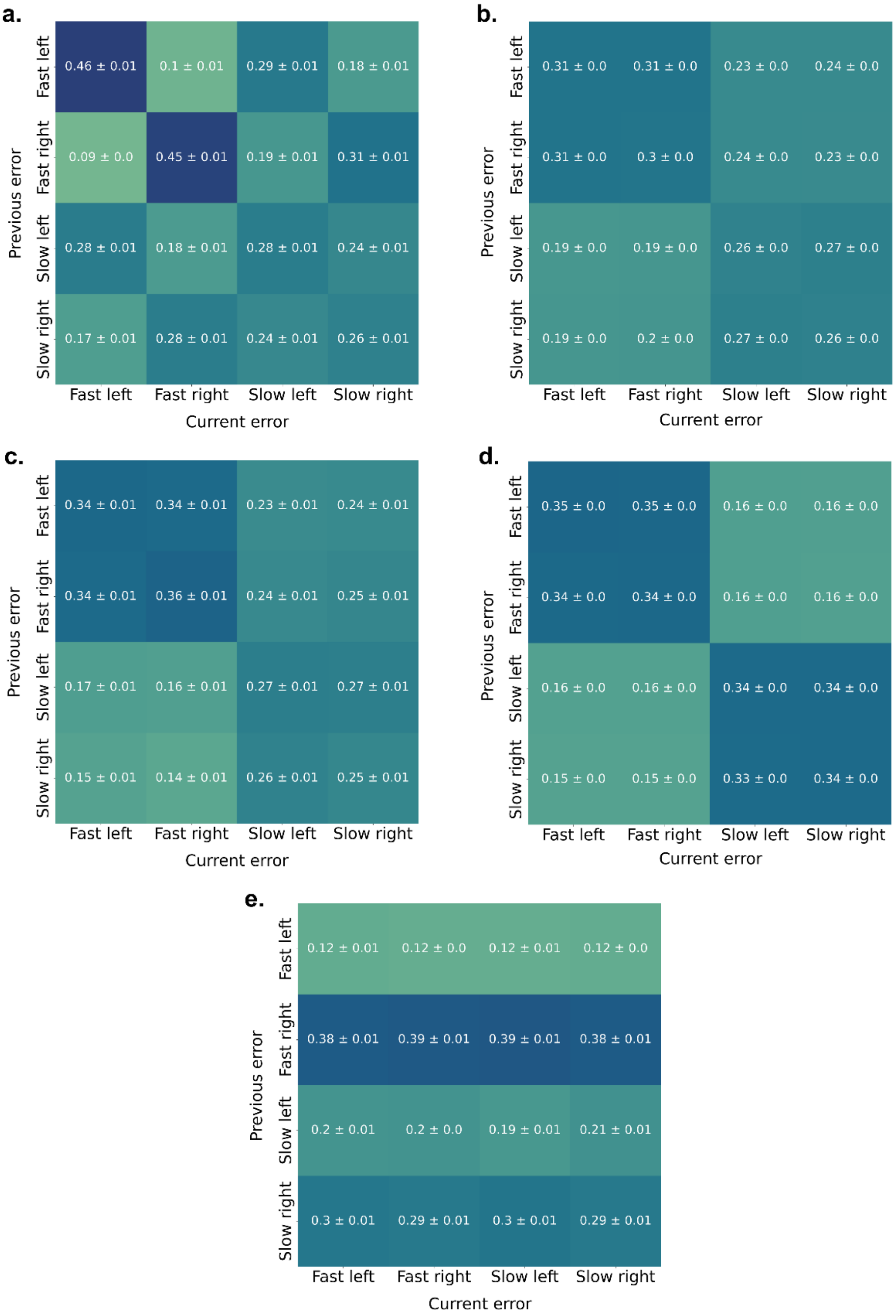


**Supplementary Figure 4: Repetition matrices for simulated data under parameter fluctuations governed by a first-order autoregressive process. a.** Simulations with time-varying starting point. **b.** Simulations with time-varying drift rate. **c.** Simulations with time-varying boundary. **d.** Simulations with fluctuations in both drift rate and boundary. **e.** Simulations with a fixed (non-fluctuating) starting point that is biased toward the 'right' boundary.

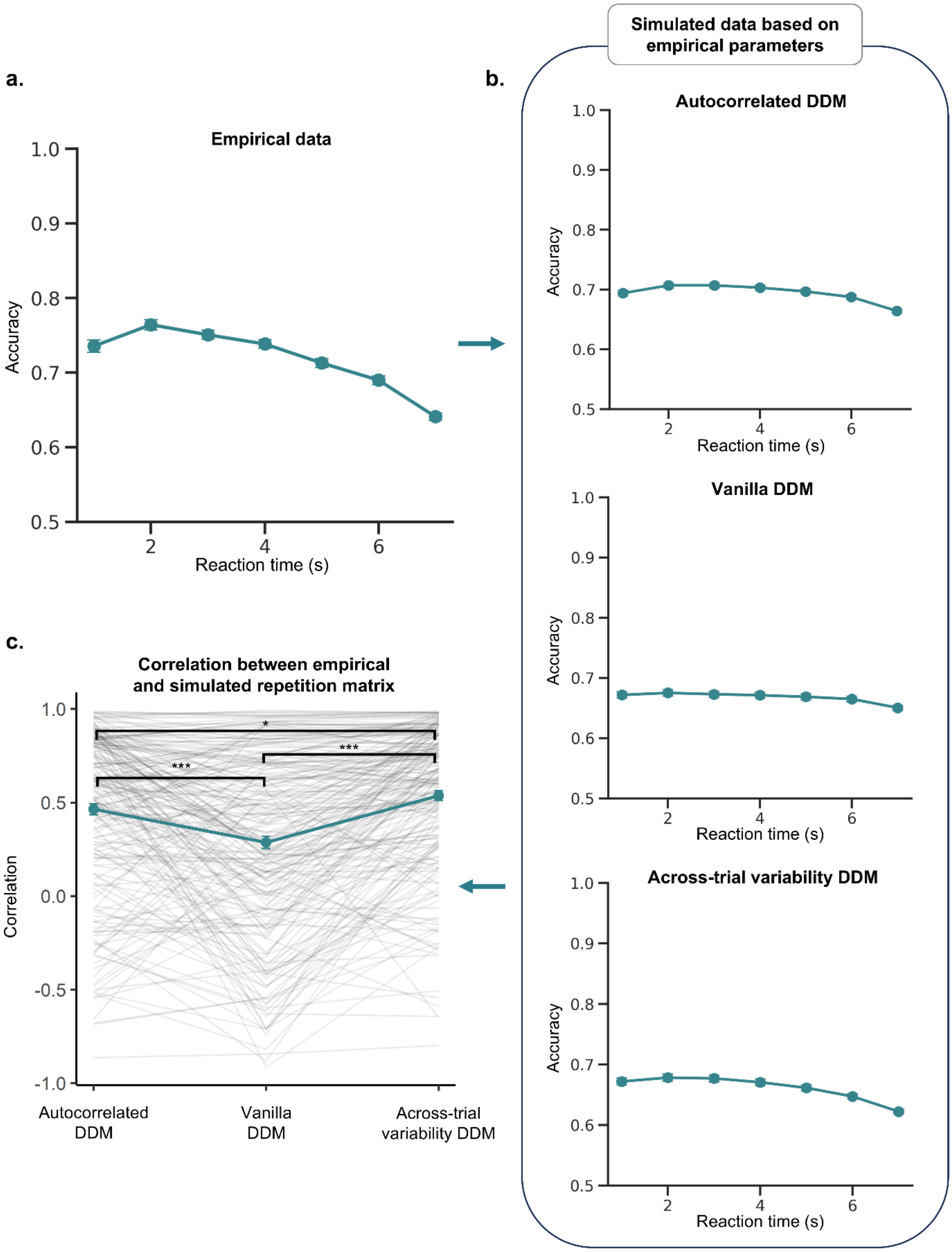


**Supplementary Figure 5: Model comparison for the conditional accuracy function (CAF) across all eight datasets. a.** The empirical CAF averaged over all 223 participants. **b.** The CAF calculated in data simulated with parameter values drawn from the posterior distribution for each of the three DDMs. The simulated CAFs for both the across-trial variability DDM and autocorrelated DDM show the presence of slow and fast errors, although less pronounced as in the empirical data. It should be noted that the vanilla DDM also seems to reproduce slow errors. However, this is presumably due to different levels of stimulus difficulty, with higher difficulty leading to both longer reaction times and reduced accuracy. **c.** Per participant, the empirical CAF was correlated with the CAF simulated with their parameter values sampled from the posterior distribution for the three DDMs. Each line represents the correlation for a participant. Linear mixed models across all datasets reveal a significant difference in correlations between the three DDMs ($F(2, 444) = 39.032$, $p < .001$, $\eta_p^2 = 0.15$). All post-hoc contrasts were significant. The correlation for the autocorrelated DDM was significantly higher than the vanilla DDM ($\beta = 0.177$, SE = 0.029, $t(444) = 6.089$, $p < .001$, $d = 0.577$, CI(95) = [0.109, 0.250]). The across-trial variability DDM showed significantly higher correlations compared to both the vanilla DDM ($\beta = 0.250$, SE = 0.029, $t(444) = 8.589$, $p < .001$, $d = 0.813$, CI(95) = [0.182, 0.319]) and the autocorrelated DDM ($\beta = -0.073$, SE = 0.029, $t(444) = -2.499$, $p = .034$, $d = -0.237$, CI(95) = [-0.141, -0.004]). For 102 out of 223 participants (45.74%), the highest correlation was observed for the across-trial variability DDM. The autocorrelated DDM and vanilla DDM were the preferred model for 81 and 40 participants (36.32% and 17.94%), respectively. Note: * = $p < .05$, *** = $p < .001$.

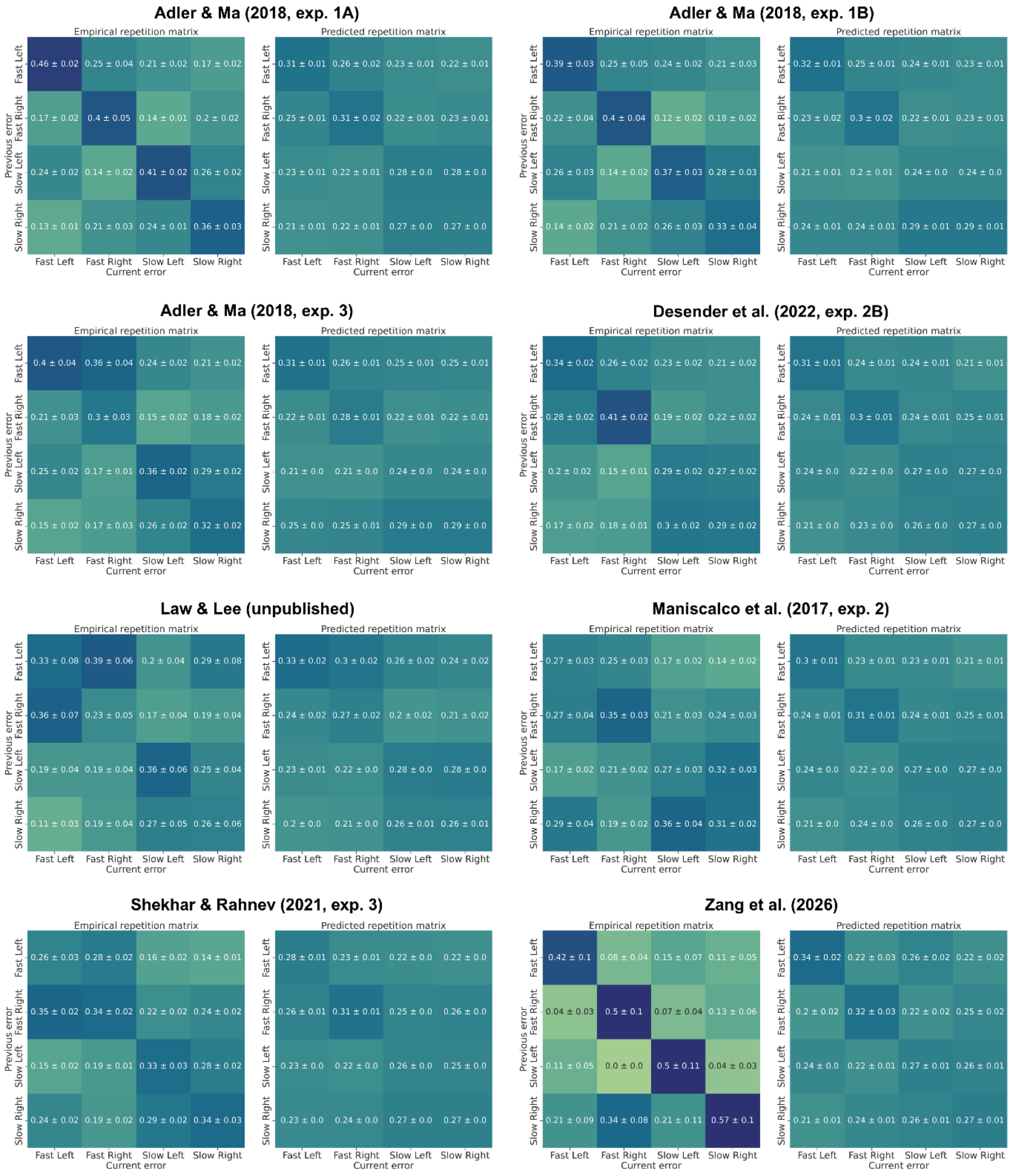


**Supplementary Figure 6: The empirical and simulated repetition matrices for each dataset separately.** Both the empirical and simulated matrices show the average over all subjects within the study (± standard error).

**Supplementary Table 1: Detailed description of empirical datasets.**

| Study | Decision description | Stimulus strength manipulation | Number of subjects | Number of trials |
|---|---|---|---|---|
| Adler & Ma (2018, experiment 1A) | Distinguish the orientation of Gabors and ellipses drawn from two distributions with different means | Gabor contrast/ ellipse aspect ratio and orientation | 11 | 2160 |
| Adler & Ma (2018, experiment 1B) | Distinguish the orientation of Gabors and ellipses drawn from two distributions from equal mean but distinct standard deviation | Gabor contrast/ ellipse aspect ratio and orientation | 11 | 2160 |
| Adler & Ma (2018, experiment 3) | Distinguish the orientation of Gabors drawn from two distributions from equal mean but distinct standard deviation | Contrast and orientation | 14 | 3240 |
| Desender et al. (2022, experiment 2B) | Decide which of two groups of dots has more dots | Dot difference | 99 | 500 |
| Law & Lee (unpublished) | Decide whether the dots are moving left or right | Motion coherence | 14 | 1200 |
| Maniscalco et al. (2017, experiment 2) | Decide on which side of the screen a Gabor patch had appeared | Contrast | 41 | 510 |
| Shekhar & Rahnev (2021, experiment 3) | Decide on orientation of Gabor patch | Contrast and orientation | 20 | 2800 |
| Zang et al. (2026) | Decide on location of Gabor patch | Contrast | 13 | 560-1040 |

**Supplementary Table 2: Overview of the prior distributions.** In the training phase, parameter values are drawn from the prior distributions and used to generate a large number of simulated datasets. The priors are chosen such that they cover a wide range of empirically plausible parameter values.

| Parameter | Prior distribution |
|---|---|
| Drift rate $v$ | $U$(0, 5) |
| Starting point $z$ | $U$(-2, 2) |
| Boundary $a$ | $U$(0.1, 5) |
| Non-decision time $T_{er}$ | $U$(0.01, 1) |
| Across-trial variability for drift rate $s_v$ | $U$(0.0, 2.5) |
| Across-trial variability for starting point $s_z$ | $U$(0.0, 2.5) |
| Autoregressive coefficient drift rate $\phi_\gamma$ | $U$(0, 1) |
| Autoregressive coefficient starting point $\phi_\zeta$ | $U$(0, 1) |
| Autoregressive coefficient boundary $\phi_\alpha$ | $U$(0, 1) |
| Error variance drift rate $\sigma^2_\gamma$ | $U$(0, 16) |
| Error variance starting point $\sigma^2_\zeta$ | $U$(0, 4) |
| Error variance boundary $\sigma^2_\alpha$ | $U$(0, 4) |
